# cONcat: Computational reconstruction of concatenated fragments from long Oxford Nanopore reads

**DOI:** 10.1101/2025.03.05.641699

**Authors:** Alexander J. Petri, Mai Thi-Huyen Nguyen, Anjali Rajwar, Erik Benson, Kristoffer Sahlin

## Abstract

Synthetic combinatorial DNA libraries are widely used to produce protein variants, optimize binders, and for high throughput studies of protein - DNA interactions. The libraries can be made by researchers or vendors and high-throughput sequencing is used for both quality control and to study the outcome of selection experiments. Oxford nanopore sequencing (ONT) is well suited to this as it allows for long read lengths and can be done rapidly with low-cost instrumentation. However, it suffers from a lower overall read accuracy and an uneven error profile. No current bioinformatics tools are well suited to the challenge of deducing the composition and order of constituent members of combinatorial libraries from ONT reads.

We introduce cONcat, an algorithm to identify the makeup of concatenated DNA fragments in a set of ONT sequencing reads from a pool of known fragments. cONcat uses the edit distance-based recursive covering algorithm for finding the best possible matchings between the fragments and the reads. In our experiments on simulated and experimental data, cONcat could accurately detect the correct fragment coverings given the short fragment sizes (< 20bp) and the sequencing errors present in ONT reads. However, we find that the high error rates in the start of ONT reads make it challenging to get confident coverage there, inferring a need for experimental strategies to avoid key sequence information in the start of reads.

## Introduction

Synthetic DNA is essential for fields such as biotechnology (Chu et al., 2024), synthetic biology (Tang et al., 2021), DNA nanotechnology (Rothemund, 2006), and DNA data storage (Ceze et al., 2019). Phosphoramidite synthesis can produce sequence-controlled oligonucleotides that can be combined in gene synthesis to produce synthetic genes up to thousands of nucleotides long (Yin et al. 2025). Although the cost of *de-novo* DNA synthesis has decreased significantly over time (Hughes & Ellington, 2017), it can be cost-prohibitive to synthesize individual gene fragments for applications where many sequence combinations are needed, such as phage display (Jaroszewicz et al., 2022), protein evolution (Stemmer, 1994), and high throughput biophysical assays (Nutiu et al., 2011). An alternative route is to combine the synthesis of short fragments with enzymatic ligation (Roth et al., 2014), Gibson assembly (Gibson et al., 2009), or primer extension (Stemmer et al., 1995) to combinatorially produce vast libraries of sequence variants from a small set of constituent strands. This produces pools of sequences with mixed identities that can be combined with selection experiments or high throughput screening to evaluate the performance of the variant libraries. In this context, high-throughput sequencing can be used to both characterize and quality control the initial library and to deduce the identity of synthetic genes that perform the selected task well. Oxford Nanopore Technologies (ONT) sequencing is compatible with libraries of diverse lengths, can be done rapidly with low-cost instrumentation, and has flow cells of various sizes. However, nanopore sequencing suffers from comparatively low sequencing accuracy that is uneven throughout the read (Ono, 2022; Delahaye & Nicolas, 2021), as we also observed in this study. The bioinformatic challenge here is to deduce the combination and order of the fragments that make up each read from a pool of known sequence fragments in an error-prone DNA sequence from Oxford nanopore data. A fragment can be missing, occur once, or several times within each read.

While algorithms capable of mapping reads to references for short and long-read sequencing data exist (Li & Durbin, 2009, Langmead et al., 2009, Sahlin, 2022, Li, 2018, Jain et al., 2018), they are not designed for our problem. These mapping tools use k-mer anchors (usually between 14 and 25 nt) between reads and the reference to guide the mapping. However, fragments can be shorter than 20nt, which in combination with ONT errors, may destroy all anchors between the fragment and the read. Since the fragments within the reads could be thought of as exons, it is tempting to apply long or short RNA-seq analysis tools, such as splice-aware mappers (Dobin, 2013), or transcript clustering (Sahlin & Medvedev, 2019, Marchet et al., 2018, Petri & Sahlin, 2024), error correction (Sahlin & Medvedev, 2021), or reconstruction tools (Petri & Sahlin, 2023, de la Rubia et al., 2022, Nip et al., 2023) to group and reconstruct the reads according to their fragment tiling make-up (mimicking distinct isoforms). However, since fragments are very short and can occur several times within a read, the data is noisy, and we know all fragments a priori, we could use this information to better find the true fragment makeup of each read than RNA tools designed for de novo reconstruction. For example, a de Bruijn graph-based assembly tool would need a large enough k-mer size to span repeats (i.e., fragments) but short enough to have shared nodes (k-mers) in the graph. Furthermore, none of the above-mentioned tools aim to produce the best tiling of fragments across the reads to find the fragment composition, which is our goal. This motivated us to design a novel algorithm tailored to our specific computational problem.

Here, we present cONcat, an algorithm that, given a pool of known fragments, finds the best fragment covering of an ONT read. Our algorithm includes a tailored edit-distance-based mapping procedure of fragment to reads, as well as an iterative fragment-selecting procedure based on the current best fragment matching a region not yet covered by a fragment. We test the algorithm’s performance using simulated data at varying read qualities and on experimentally produced data from a set of DNA products randomly ligated from a library of sequence fragments of varying lengths that are sequenced with ONT.

## Methods

### Preliminaries

Let *r* ∈ *R* denote a string consisting of letters in Σ = (*A, C, G, T*), that represents a sequencing *read*. Similarly, let *f* ∈ F denote a string of alphabet Σ representing a sequence fragment. Let *r*[*i, j*] indicate the substring starting at *i* and ending at *j* in *r*. We let | · | denote the length of a string. We align the fragments to *r*. Let *E*(*f, r*) denote the *edit distance* between *f* and *r* after a semi-global alignment of *f* to *r* (i.e., gaps at the start and end of *f* are not penalized). The edit distance is the minimum number of single-character edits (insertions, deletions, substitutions) between two strings required to transform a string into another string. We refer to the *covering* of a read as the process of, through pairwise alignment between the fragments and a read, finding the locations of fragments in the read. We say that a read is fully covered if all the correct fragments that make up the read have been identified. In simulated data, we know if a read is fully covered or not, while it is not always possible in experimental data.

### Algorithm

Our algorithm attempts to find the best non-overlapping covering of fragments on *r* and works greedily. All fragments are aligned to the read, and the best matching location is identified using the alignment identity. Let *A*(*f*_*i*_, *r*) denote an alignment produced from the semi-global edit distance computation between a fragment *f*_*i*_ and *r* (ignoring the gaps at ends). We then compute the alignment identity *I*(·) of *f*_*i*_ to *r* as 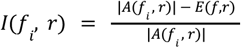.

When the fragment location (say, aligned to *r*[*i, j*]) with the best alignment identity has been identified, we assign the fragment to that location and split the read into two, namely *r*[0, *i*] and *r*[*j*, |*r*|]. We then treat the two new subsequences of *r* as new reads and recursively identify the best fragment matches within the two sub-reads and split them. Should two fragments have the same alignment identity, we choose the first fragment with the highest alignment identity.

The algorithm terminates as soon as no sub-sequence longer than 5 base pairs has yet to be mapped with fragments or if all remaining sub-sequences cannot be mapped with fragments due to their alignment identity being below a minimum identity threshold *T* (set to 0.75). Should the alignment identities of all fragments be below *T*, the sequence remains unmapped.

### Input, output, and implementation details

Our algorithm’s input consists of base-called reads generated via ONT sequencing in the fastq format. The algorithm outputs two CSV files. One of the CSV files contains the position and edit distance of a detected fragment in a read, where a single read occupies multiple lines (one for each fragment). The second CSV file shows the percentage of bases per read the algorithm covered with fragments. The algorithm was implemented in the Rust programming language and is available via https://github.com/aljpetri/cONcat. We use rust-bio library (Köster) to parse the reads and edlib-rs (Both), a Rust portation of the edlib algorithm (Šošić and Šikić) to estimate the edit distances between the fragments and the reads. Edlib can be sped up by setting a maximum allowed edit distance. We use our alignment identity threshold *T* and limit the maximum edit distance *k* for which hits are found by calculating *k* = *ceil*(|*f*|(1/1 − *T*)).

### Generating fragment libraries

The simulated and experimental datasets are produced from constituent sequence fragments initially generated from a Python script that produced a set of 20 nt long sequences that have no homo-polymers, similar melting temperatures and a high degree of sequence orthogonality to the other members of the set. This set was used to produce linear DNA fragments of varying lengths flanked by constant regions to form non-palindromic sticky ends in both ends. We designed two such fragments of each length between 10-20 nt long (22 fragments in total). We also designed two ‘capping’ fragments that form one sticky end and one blunt end.

### Experimental methods

We developed an experimental dataset based on randomly ligated DNA fragments. The oligonucleotides needed to form the sequence fragments were synthesized by ‘Integrated DNA Technologies (IDT) with 5’ phosphate groups added to all strands except for the strands forming the 5’ end of the capping fragments. The forward and backward strands comprising each fragment were mixed separately at 10 uM each in 1x T4 ligation buffer (New England Biolabs). These mixtures were annealed using a 30-minute temperature ramp from 80 C to 20 C in a thermal cycler. The annealed fragments were then mixed in a single tube and a T4 ligase enzyme was added (New England Biolabs), this was incubated at 16 C for 10h followed by an overnight incubation at 4 C to fuse the fragments (Fig. 2A). The ligated products were purified using Ampure XP beads (Beckman Coulter) to remove unligated products and enzymes/reaction buffer. This was used as a template for a PCR reaction with primers targeting the cap fragments that feature at the start and end of many ligated products. This PCR reaction was again purified using Ampure XP beads to remove excess primers and PCR buffer. After this, the Oxford nanopore ligation sequencing kit (LSK-114) was used to add sequencing adapters to the library, with the supplied DNA control sample (DCS) included. This was sequenced on a R10.4.1 flongle flow cell using a minion sequencer. Super accurate base calling was used in Minknow. For details on the DNA amplification, AMPure Purification Protocol, and ONT sequencing, see Supplementary materials.

**Figure 1.**
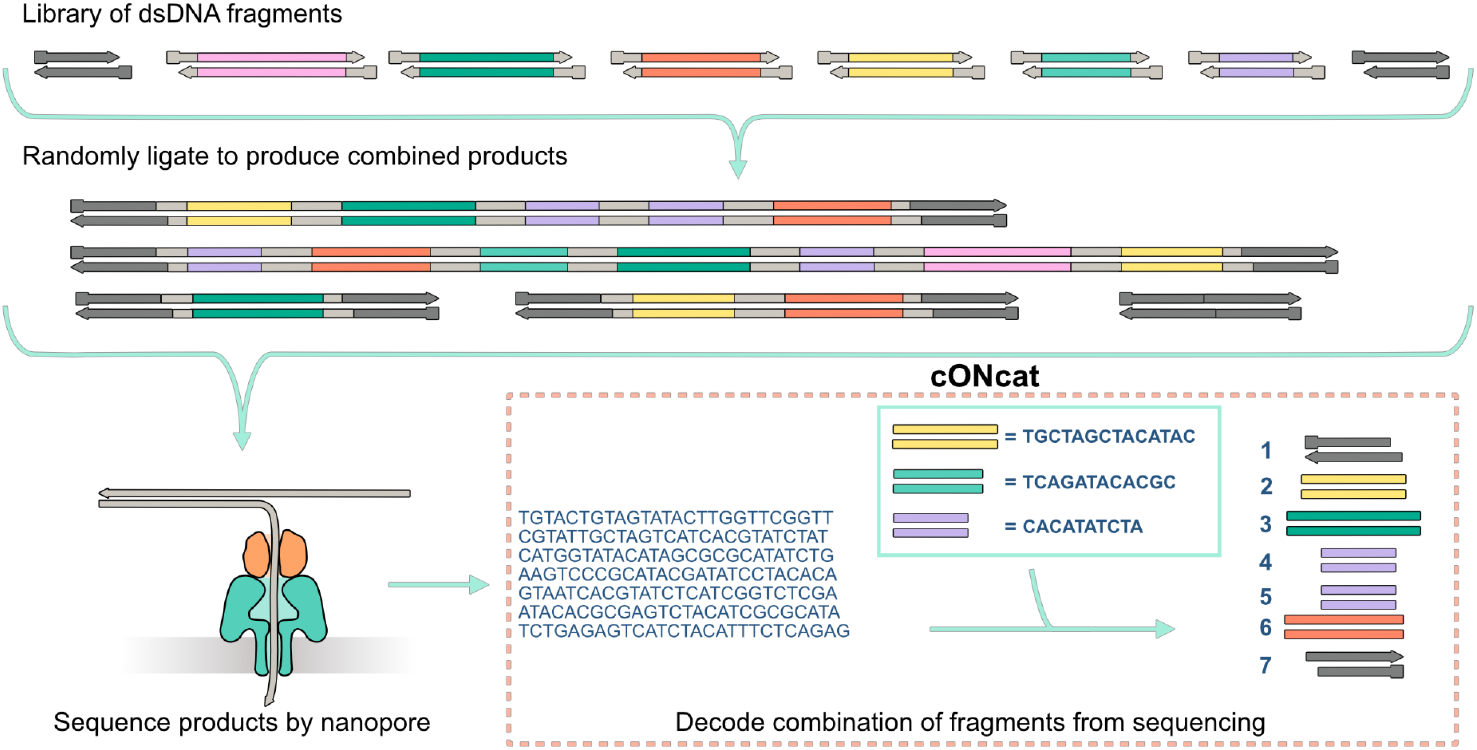
An overview of the experimental workflow. DNA fragments are randomly concatenated by ligation to form a pool of longer products. The products are sequenced using Oxford nanopore sequencing to generate basecalled reads. The reads are input together with the list of initial fragments in cONcat to generate an ordered list of the constituent fragments in each read.

**Figure 2.**
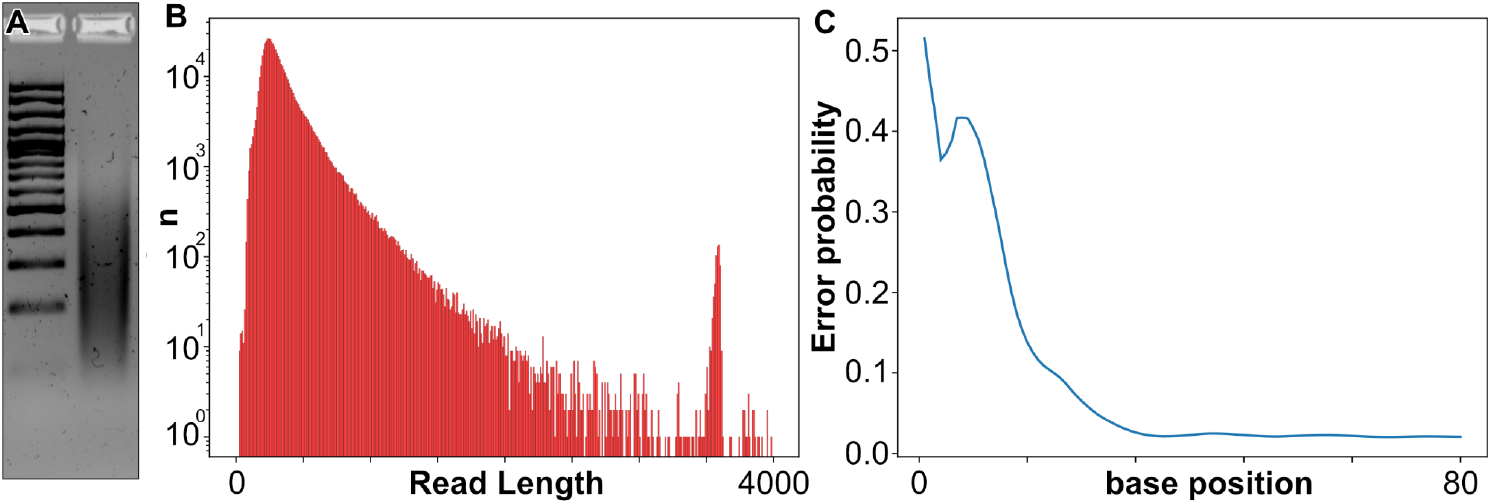
Experimental assembly and sequencing of a library dataset. (**A**) Agarose gel electrophoresis on ligated products (right) compared to 100 bp generuler plus ladder (left). (**B)** Histogram of read lengths from nanopore sequencing. (**C**) Average error probability per base in the first 80 read bases from nanopore sequencing.

The sequencing resulted in 559,080 reads passing the quality filter with a mean read length of 353 nt, a smooth distribution of read lengths we present with a spike in read length at around 3600 nt, corresponding to the nanopore control sample (Fig. 2B). The average error probability was calculated from the nanopore Phred quality score (Fig. 2C). It revealed a very high error rate in the first 10 nucleotides of around 40% or more, followed by a rapid decrease toward around 15% at nucleotide 20. The error rate then stabilized at around 2% after nucleotide 40. This indicates that the error profile is drastically different at the start of the read compared to the middle, and the chance of successfully aligning fragments may be quite different.

### Simulating reads

We simulated datasets with varying error rates and error profiles. First, we simulated sets of reads with uniform error rates over the full length of the reads. We used the set of 22 fragments that we used to synthesize biological data. In the simulations, we concatenated ten fragments per read (randomly drawn with replacement). We then applied the errors to the reads. The error rates (in percent) we used for these experiments are 0, 1, 5, 7, 10,15, 20, 25, and 30, and we denote them as SIM0, …, SIM30, respectively.

However, ONT reads are known to contain non-uniform error rates with high error rates in the start and end regions (as mentioned in (Ono, 2022)) with the middle part containing relatively low error rates. Therefore, we additionally simulated a dataset SIM_NU (for Non-Uniform errors) with a higher error rate at the beginning of the reads (Fig. 2C). Using our experimental data from real ONT sequencing reads, we calculated an average per-base error rate of the first 100 nucleotides from 557k ONT reads and used the average error rates per base to simulate the SIM_NU dataset of 1000 reads with the non-uniform error profiles accordingly. Specifically, the first 100 bases in the simulated read have individual per-base error rates as computed from experimental data, and the remainder of the read is simulated with the average error rate computed from position 51 to 100.

We additionally simulated a dataset (SIM_Rand) consisting of 1000 reads, each consisting of 1000nt simulated uniformly at random to assess the false discovery rates of our algorithm for different alignment identity thresholds *T*. As we know the ground truth for all our simulated datasets, we can calculate the percentage of true and falsely detected fragments in each read.

## Results

### Simulated datasets with uniform error rates

We tested cONcat on SIM0 -- SIM30 and with different settings of threshold *T*. Figure 3 shows the number of reads that could fully be covered with fragments by cONcat for different error rates and settings of T. While our algorithm can correctly cover all reads for low error rates of 0 to 3 percent, higher error rates yield incorrect coverings. As expected, higher settings of *T* (more conservative mapping), produce fewer full coverings. Figure 4 shows the number of correctly (green) and incorrectly (red) mapped fragments for each error rate and setting of *T*. cONcat places nearly all fragments correctly up to 10% error rate, suggesting robustness against errors.

**Figure 3.**
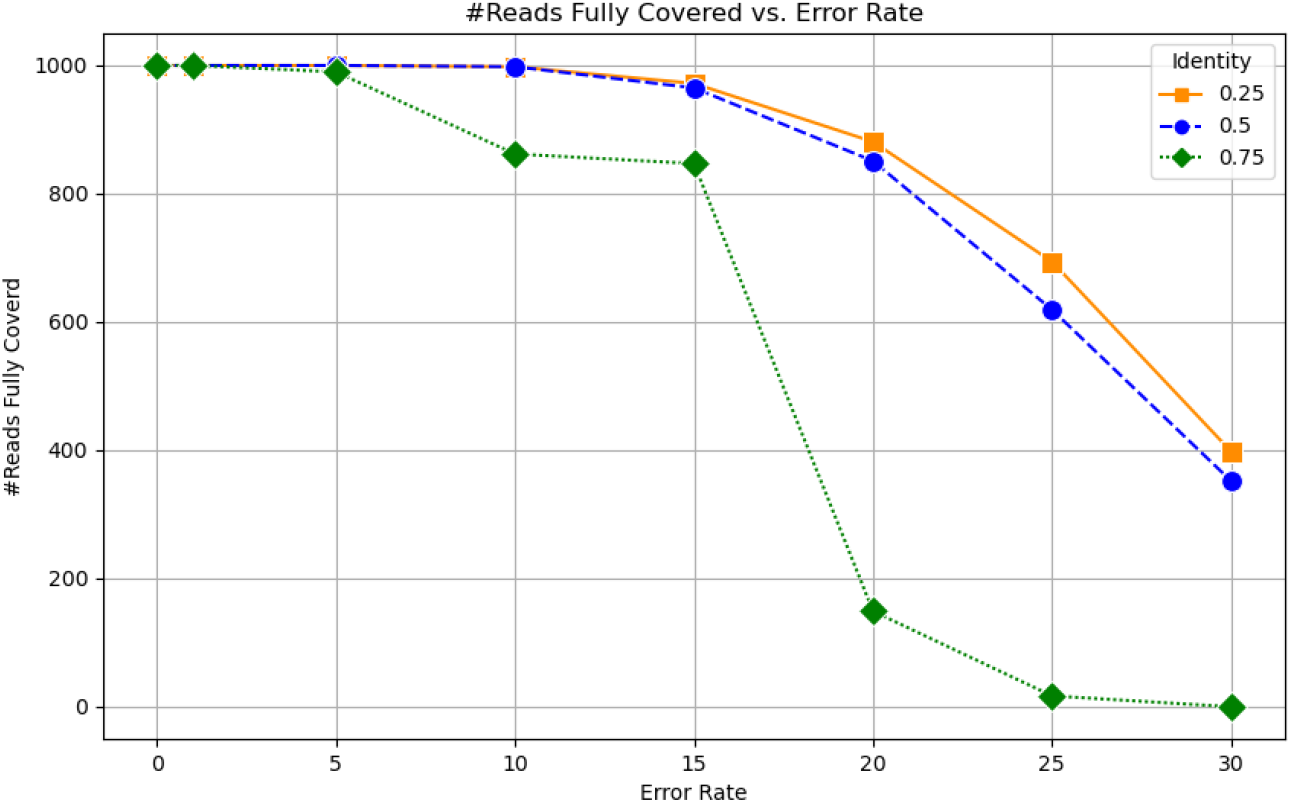
Number of reads fully mapped for different error rates and settings of *T* (identity). While for low error rates, all settings yield fully correct mappings of all reads the correctness of the mappings decreases with increasing error rates. Using lower settings of *T* results in higher rates of correct mappings.

**Figure 4.**
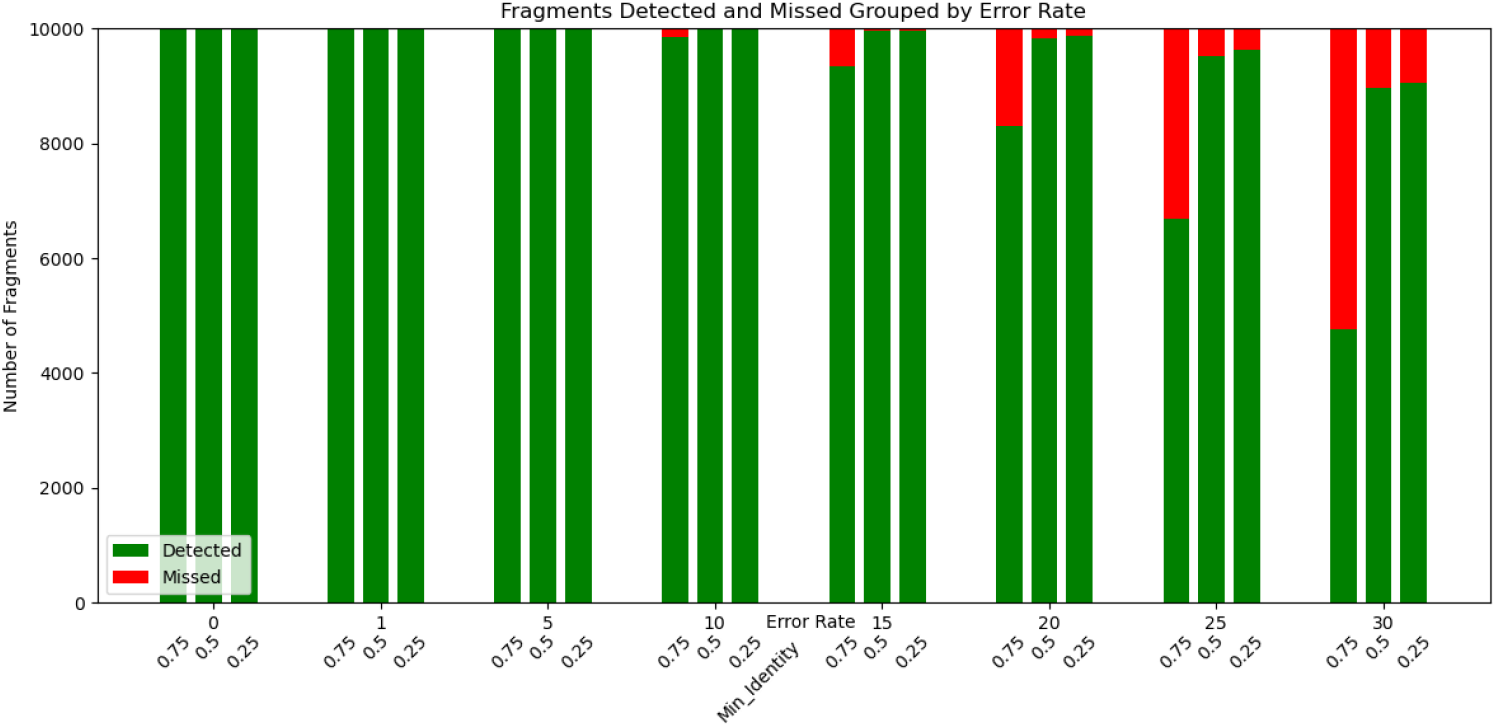
Number of fragments correctly mapped (green) and incorrectly mapped (red) for each error rate and different identity settings (0.25, 0.5, and 0.75) of *T*.

### Simulated datasets with non-uniform error rates

We also tested the cONcat algorithm with different thresholds *T* on SIM_NU with a more challenging error profile at the beginning of the reads. For *T* = 0. 75, cONcat fully covered only 249 (24.9%) out of 1000 reads correctly. As expected, the beginnings of reads with a high error rate were typically incorrectly or not covered for most of the reads that were not fully correctly covered. Out of the 10,000 possible fragment locations in the 1,000 reads, 781 fragments were missed with *T* = 0. 75. These 781 fragments were mostly residing as the first fragment in the 751 reads that were not fully covered. When lowering the identity threshold *T* to 0.5, 629 reads (62.9%) were fully and correctly covered. Out of the 10,000 possible fragment locations in the 1,000 reads, 378 fragments were missed with *T* = 0. 5. Similar to the setting *T* = 0. 75, the large majority of the 371 reads that were not fully covered had one missing fragment in the beginning.

### False fragment discoveries

While a lower value of T yielded more correct fragment coverings (i.e., higher sensitivity), it can make incorrect assignments by taking the best matching fragment that happened to have an alignment identity above the threshold (false positive). We used our fully random dataset SIM_Rand to assess the number of false positive fragment mappings for different values of *T*. For this dataset we do not want any of our fragments to map to the reads, as they are all false positives. We found that the number of false positives increases when lowering *T*. For *T* = 0. 75, the algorithm mapped 43 fragments to the full dataset, while it mapped 29,655 and 37,534 fragments to the data when *T* were set to 0.5 and 0.25, respectively. This indicates that if our DNA products consist of sequences other than our fragments, they could potentially be covered with fragments by chance when setting *T* to 0.5 or lower.

### Experimental data

We then applied our algorithm to the sequenced data using different values of the minimum identity thresholds, from 0.5 to 0.75, and studied the effect of alignments in the start of reads. With a threshold of 0.75, cONcat typically only maps fragments after 30-40 nt into the reads (Fig. 5), meaning that the first one or two fragments of the reads become unmapped. By decreasing the threshold, this effect is reduced, and at a threshold of 0.5, most reads have their first alignment starting before nucleotide 10. When looking at the coverage fraction of reads, we note that most reads have a coverage over 70% irrespective of the alignment threshold *T* (Fig. 5) although, as expected, the coverage is higher for lower thresholds. A clear outlier here is the nanopore control sample, where we do not expect any matches (similarly to the SIM_Rand dataset). The control sample forms a clear spot at read length 3600 nt. Interestingly, this spot has a very low coverage at threshold 0.75, as is expected, since it is not composed of the fragments in question. When the threshold is reduced, the control sample increases its coverage since more fragments of the library can be aligned to reads when more mismatches are allowed, forming a false positive. As the threshold is modified, the coverage of the non-control portion of the reads also changes. When the threshold decreases, more alignments are found in the ends of reads, increasing the overall coverage. This effect is most pronounced in shorter reads where the ends make up a larger fraction leading to a change in the overall distribution (Fig. 5).

**Figure 5.**
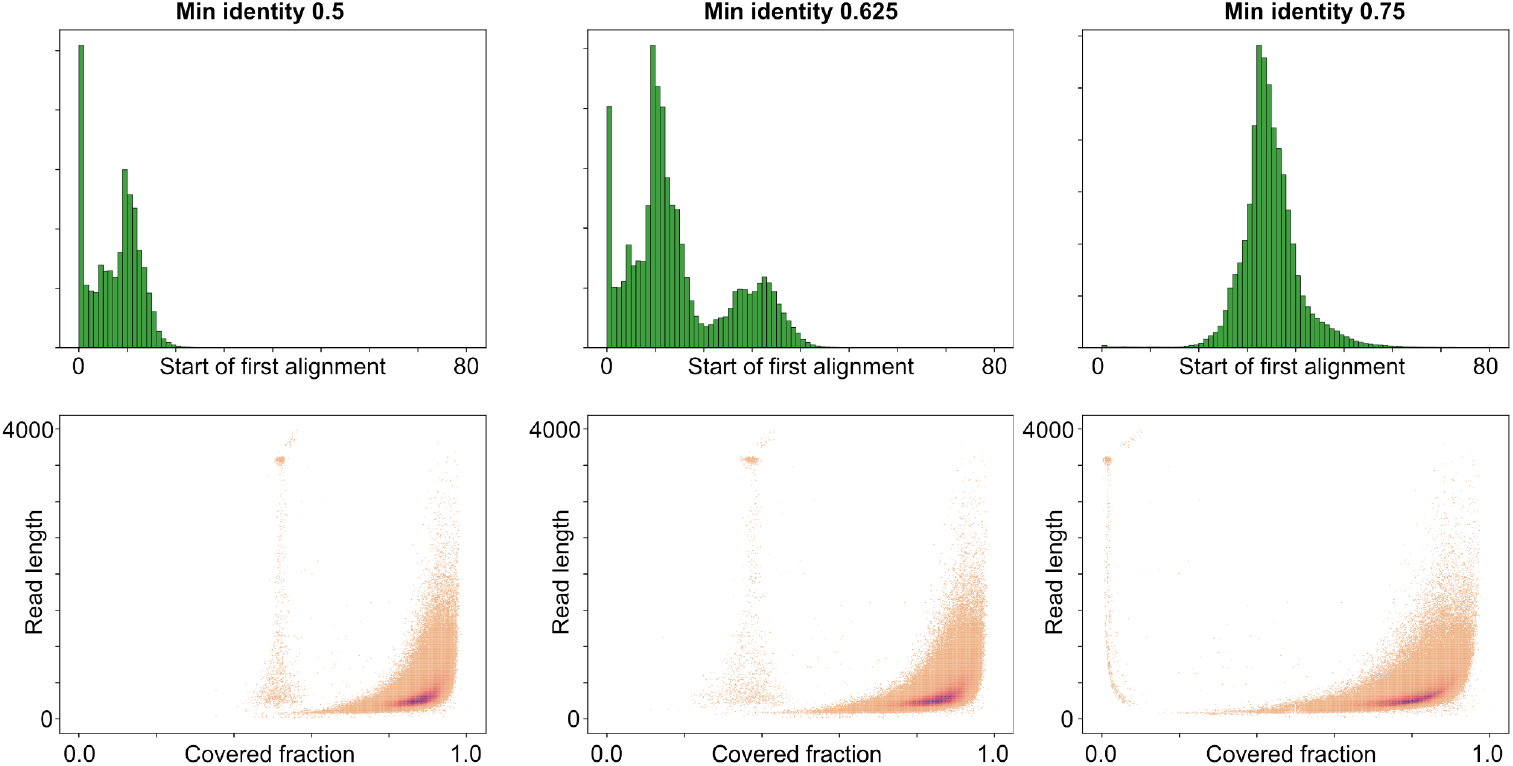
Processing of sequenced data with varying minimum alignment identity threshold *T*. Top: Histogram of start position of first alignment found in each read. Bottom: 2D histogram of the fraction of read coverage versus read length. The nanopore control samples formed an island at read length 3600 nt.

### Memory and time

On our data, cONcat has a low memory consumption and runtime. Our algorithm processes a simulated dataset of 1000 reads in 6 seconds (Suppl. Fig. 1) or less using less than 6Mb (Suppl. Fig. 2) of RAM. On our experimental data consisting of 559,080 reads, cONcat processes this dataset in 2,309 seconds using 949 Mb of RAM.

## Discussion

Today, synthetic combinatorial DNA libraries are widely used to produce protein variants, optimize binders, and for high throughput studies of protein - DNA interactions. Libraries can be synthesized by commercial vendors or directly by researchers using techniques such as ligation, Gibson assembly or primer extension. High throughput DNA sequencing allows for the control of quality, diversity and homogeneity of the libraries before experiments, and can be used again in selection experiments to validate what sequence variants were favoured by the selection pressures. In this context, Oxford nanopore sequence has several attractive properties: it allows for long reads, it is rapid, and it can be done with inexpensive instrumentation. Although the read quality of Oxford nanopore sequencing has steadily increased with improved chemistry and base calling algorithms, it has significantly lower accuracy than Illumina or PacBio sequencing, and the error profile is not homogeneous throughout the reads.

We introduced cONcat, an algorithm to identify the fragment composition in Synthetic DNA products that have been sequenced with ONT sequencing reads. Unlike other read mapping or sequence reconstruction tools (such as genome or transcriptome assembly tools), cONcat is designed specifically for the computational problem of finding a covering of fragments across the reads with as high alignment identity as possible. cONcat utilizes edit distance alignment to find the best-fitting local fragment to iteratively assign fragments to the sequencing reads.

To assess the performance of our algorithm, we tested the tool on simulated and experimental datasets. Using simulated data with a random uniform error profile at various error rates (SIM0-SIM30), we showed that cONcat can accurately identify the correct fragments at error rates up to around 10-15% errors, despite the fact that fragments can be shorter than 20nt. Our algorithm uses a threshold T as a minimum cutoff identity to prevent overfitting fragments to the reads. We tested our algorithm with different settings of T on mapping fragments to random DNA sequences (SIM_Rand) to assess false positive fragment identification rates, and a threshold at 0.5 or lower resulted in many false positives. We further assessed the performance of our algorithm on reads with non-uniform error profiles (SIM_NU), as observed in our experimental data. Our experiments show that with higher error rates in the beginning of reads, cONcat is able to correctly place much fewer fragments at the beginning of the reads, which is expected. Furthermore, cONcat has a low resource usage with the runtime slightly increasing with lower settings of *T*. Since the algorithm can be easily parallelized per read, it can handle much larger datasets within a reasonable time if needed.

For our experimental data, we observe that cONcat with *T = 0*.*75* typically only maps fragments after 30-40 nt into the reads (Fig. 5) due to the high error rate. However, most reads are covered over 70% with fragments. Furthermore, the control sample is clearly distinguished from our synthetic DNA products as they typically have a coverage of around 0-5% (Fig. 5) with the stringent threshold T=0.75.

### Future work

Reducing the threshold gradually increases the amount of alignments found in the start of the reads, although it simultaneously increases the coverage of the control sample, indicating more false positive alignment. In applications where correct mapping of fragments in the start of reads is crucial, experimental strategies such as PCR primer extension that extend the length of the library may be needed to overcome the very high error rate in the start of ONT reads by moving the real library further into the read.

As we observed, the regions with higher error rates within reads are not correctly covered by cONcat. This is mainly due to incorrect base calls that yield sequences too dissimilar to the fragments. A possible future work could therefore include an additional, more carefully tailored covering step for sequences that cONcat could not cover with the default alignment identity parameter *T*. Such a step could, e.g., utilize the Phred quality values within the read by taking them into account when performing the pairwise alignment between the fragments and the read.

## Data availability

The algorithm was implemented in the Rust programming language and is available via https://github.com/aljpetri/cONcat. The same repository also contains all scripts to simulate the data used for evaluation in the paper. The experimental data is available at figshare with DOI: https://doi.org/10.6084/m9.figshare.28524599.v2.

## Acknowledgments

Kristoffer Sahlin was supported by the Swedish Research Council (SRC, Vetenskapsrådet) under Grant No. 2021-04000. Erik Benson was supported by the Swedish Research Council (SRC, Vetenskapsrådet) under Grant No.2022-0414. The computations were enabled by resources provided by the National Academic Infrastructure for Supercomputing in Sweden (NAISS), partially funded by the Swedish Research Council through grant agreement no. 2022-06725.

## Bibliography

Both, jean-pierre. 2020. “edlib_rs.” github edlib-rs. https://github.com/jean-pierreBoth/edlib-rs.

Ceze, Luis, Jeff Nivala, and Karin Strauss. 2019. “Molecular digital data storage using DNA.” Nature Reviews Genetics 20 (8): 456–466. 10.1038/s41576-019-0125-3.

Chu, Alexander E., Tianyu Lu, and Huang Po-Ssu. 2024. “Sparks of function by de novo protein design.” Nature Biotechnology 42 (2): 203–215. 10.1038/s41587-024-02133-2.

Delahaye, Clara, and Jacques Nicolas. 2021. “Sequencing DNA with nanopores: Troubles and biases.” PloS one 16 (10).

de la Rubia, Ivan, A. Srivastava, W. Xue, J. A. Indy, S. Carbonell-Sala, J. Lagarde, M. Mar Albà, and Eduardo Eyras. 2022. “RATTLE: reference-free reconstruction and quantification of transcriptomes from Nanopore sequencing.” Genome Biology 23 (1): 153.

Dobin, Alexander, Carrie A. Davis, Felix Schlesinger, Jorg Drenkow, Chris Zaleski, Sonali Jha, Philippe Batut, Mark Chaisson, and Thomas R. Gingeras. 2013. “STAR: ultrafast universal RNA-seq aligner.” Bioinformatics 29 (1): 15–21.

Gibson, Daniel G., Lei Young, Ray-Yuan Chuang, J Craig Venter, Clyde A. Hutchinson III, and Hamilton O. Smith. 2009. “Enzymatic assembly of DNA molecules up to several hundred kilobases.” Nature Methods 6 (5): 343–345. 10.1038/nmeth.1318.

Hughes, Randall A., and Andrew D. Ellington. 2017. “Synthetic DNA Synthesis and Assembly: Putting the Synthetic in Synthetic Biology.” Cold Spring Harb Perspect Biol 9 (1): a023812. 10.1101/cshperspect.a023812.

Jain, Chirag, Alexander Dilethey, Sergey Koren, Srinivas Aluru, and Adam M. Phillippy. 2018. “A fast approximate algorithm for mapping long reads to large reference databases.” Journal of Computational Biology 25 (7): 766–779.

Jain, Chirag, Arang Rhie, Haowen Zhang, Claudia Chu, Brian P. Valenz, Sergey Koren, and Adam M. Phillippy. 2020. “Weighted minimizer sampling improves long read mapping.” Bioinformatics 36 (Supp_1): i111–i118.

Jaroszewicz, Weronika, Joanna Morcinek-Orłowska, Karolina Pierzynowska, Lidia Gaffke, and Grzegorz Węgrzyn. 2022. “Phage display and other peptide display technologies.” FEMS Microbiology Reviews 46 (2): fuab052. 10.1093/femsre/fuab052.

Köster, Johannes. 2016. “Rust-Bio: a fast and safe bioinformatics library.” Bioinformatics 32 (3): 444–446. https://academic.oup.com/bioinformatics/article/32/3/444/1743419.

Langmead, Ben, Cole Trapnell, Mihai Pop, and Steven L. Salzberg. 2009. “Ultrafast and memory-efficient alignment of short DNA sequences to the human genome.” Genome biology, no. 10, 1–10.

Levenshtein, V. I. 1966. “Binary codes capable of correcting deletions, insertions, and reversals.” Proceedings of the Soviet physics doklady 10 (8): 707–710. http://profs.sci.univr.it/∼liptak/ALBioinfo/2012_2013/2011_2012/files/levenshtein66.pdf.

Li, Heng. 2018. “Minimap2: pairwise alignment for nucleotide sequences.” Bioinformatics 34 (18): 3094–3100.

Li, Heng, and Richard Durbin. 2009. “Li, Heng, and Richard Durbin. “Fast and accurate short read alignment with Burrows–Wheeler transform.” bioinformatics 25 (14): 1754–1760.

Marchet, Camille, Lolita Lecompte, Corinne da Silva, Corinne Cruaud, Jean-Marc Aury, Jaques Nicolas, and Pierre Peterlongo. 2018. “CARNAC-LR : Clustering coefficient-based Acquisition of RNA Communities in Long Reads.” JOBIM 2018-Journées Ouvertes Biologie, Informatique et Mathématiques. 2018, 1–3.

Nip, Ka Ming, Saber Hafezqorani, Kristina K. Gagalova, Readman Chiu, Chen Yang, René L. Warren, and Inanc Birol. 2023. “Reference-free assembly of long-read transcriptome sequencing data with RNA-Bloom2.” Nature Communications 14 (1): 2940.

Nutiu, Razvan, Robin C. Friedman, Shujun Luo, Irina Khrebtukova, David Silva, Robin Li, Lu Zhang, Gary P. Schroth, and Christopher B. Burge. 2011. “Direct measurement of DNA affinity landscapes on a high-throughput sequencing instrument.” Nat Biotechnol 29 (7): 659–664. 10.1038/nbt.1882.

Ono, Yukiteru. 2022. “PBSIM3: a simulator for all types of PacBio and ONT long reads.” NAR genomics and bioinformatics 4 (4): qac092.

Petri, Alexander J., and Kristoffer Sahlin. 2023. “isONform: reference-free transcriptome reconstruction from Oxford Nanopore data.” Bioinformatics 39 (Supplement_1): i222–i231.

Petri, Alexander J., and Kristoffer Sahlin. 2024. “De novo clustering of extensive long-read transcriptome datasets with isONclust3.” bioRxiv 2024-10.

Roth, Theodore L., Ljiljana Milenkovic, and Matthew P. Scott. 2014. “A Rapid and Simple Method for DNA Engineering Using Cycled Ligation Assembly.” Plos One 9 (9): e107329. 10.1371/journal.pone.0107329.

Rothemund, Paul W. 2006. “Folding DNA to create nanoscale shapes and patterns.” Nature 440 (7082): 297–302. 10.1038/nature04586.

Sahlin, Kristoffer. 2022. “Strobealign: flexible seed size enables ultra-fast and accurate read alignment.” Genome Biology 23 (1): 260.

Sahlin, Kristoffer, and Paul Medvedev. 2019. “De novo clustering of long-read transcriptome data using a greedy, quality-value based algorithm.” Research in Computational Molecular Biology: 23rd Annual International Conference, RECOMB 2019 Proceedings 23:227–242.

Sahlin, Kristoffer, and Paul Medvedev. 2021. “Error correction enables use of Oxford Nanopore technology for reference-free transcriptome analysis.” Nature Communications 12 (1): 2.

Šošic, Martin, and Mile Šikić. 2017. “Edlib: a C/C ++ library for fast, exact sequence alignment using edit distance.” Bioinformatics 33 (9): 1394–1395. https://academic.oup.com/bioinformatics/article/33/9/1394/2964763?login=false.

Stemmer, Willhelm P., Andreas Crameri, Kim D. Ha, Thomas M. Brennan, and Herberg L. Heyneker. 1995. “Single-step assembly of a gene and entire plasmid from large numbers of oligodeoxyribonucleotides.” Gene 164 (1): 49–53. 10.1016/0378-1119(95)00511-4.

Stemmer, Willhem P. 1994. “Rapid evolution of a protein in vitro by DNA shuffling.” Nature 370 (6488): 389–391. 10.1038/370389a0.

Tang, Tzu-Chieh, Bolin An, YuanYuan Huang, Sangita Vasikaran, Yanyi Wang, Xiaoyu Jiang, Timothy K. Lu, and Chao Zhong. 2021. “Materials design by synthetic biology.” Nature Reviews Materials 6 (4): 332–350. 10.1038/s41578-020-00265-w.

Yin, Yipeng, Reed Arneson, Yinan Yuan, and Shiyue Fang. 2025. “Long oligos: direct chemical synthesis of genes with up to 1728 nucleotides.” Chemical Science 16, no. 4 (Jan): 1966--1973.

