## Supplementary file for "cONcat: Computational reconstruction of concatenated fragments from long Oxford Nanopore reads"

### Supplementary Material

#### DNA Library Preparation

We developed a diverse library of DNA fragments with varying sizes. To assemble the library, the fragments were annealed under a controlled temperature ramp, gradually cooling from 95°C to 4°C over a duration of 30 minutes. This process facilitated the precise hybridization of complementary strands.

The annealed fragments were subsequently ligated using T4 DNA ligase (New England Biolabs) at a 10h incubation at 16°C followed by an overnight incubation at 4°C, ensuring efficient covalent linkage between DNA fragments. This ligation step resulted in the creation of a heterogeneous library of double-stranded DNA (dsDNA).

The ligated DNA library was subjected to PCR amplification using primers complementary to the non-phosphorylated start and end sequences incorporated during the ligation reaction. These non-phosphorylated sequences functioned as terminal caps, preventing continuous extension of the ligated fragments and ensuring controlled amplification. A gradient PCR was performed to determine the optimal annealing temperature, with reactions conducted at 50°C, 53.6°C, 55°C, and 58°C for 15 cycles. Amplification was observed across all tested temperatures, and 58°C was selected for the final amplification to minimize non-specific primer binding. The PCR reaction setup is detailed in Table 1.

| Component | Volume (µl) |
|---|---|
| 2x DreamTaq Master Mix | 100 |
| Fwd primer-Start (10 µM) | 10 |
| Bwd primer-End (10 µM) | 10 |
| Ligated sample | 2 |
| H2O | 78 |
| Total Vol (ul) | 200 |

***Table 1: PCR Reaction Setup***

Following amplification, the PCR products were purified using AMPure XP beads to remove unreacted components or smaller by-products ensuring high-purity amplicons for downstream nanopore sequencing. Sequencing was performed following the standardized protocol provided by the Nanopore Community, using the Flonglebranch of the Ligation Sequencing Amplicons V14 (SQK-LSK114) workflow.

##### Nanopore Sequencing Workflow

##### End-Preparation of DNA

For the end-prep reaction, 100 fmol of purified, ligated DNA was required. The DNA concentration was precisely measured using a Qubit fluorometer, assuming an average amplicon length of 0.5 kbp. Based on this estimation, 32.5 ng of DNA was used to achieve the required input amount. The sample was diluted in nuclease-free water to a final volume of 24.5 µL before proceeding with the reaction setup, which was performed according to the recommended protocol (Table 1).

| Reagent | Volume (µL) |
|---|---|
| DCS | 0.5 |
| DNA | 24.5 |
| Ultra II End-Prep Reaction Buffer | 3.5 |
| Ultra II End-Prep Enzyme Mix | 1.5 |
| Total | 30 |

***Table 2: End-Prep Reaction Setup***

The reaction was incubated in a thermocycler at 20°C for 5 minutes, followed by 65°C for 5 minutes to complete the end-prep process. Upon completion, the sample was purified using AMPure XP beads and transferred to a clean 1.5 mL Eppendorf DNA LoBind tube for downstream processing.

##### AMPure Purification Protocol

1. **Bead Binding:** Add 30 µL of resuspended AMPure XP beads (AXP) to the end-prep reaction and mix gently by flicking the tube.
2. **Incubation:** Place the tube on a Hula mixer at room temperature for 5 minutes to allow DNA binding.
3. **Washing Steps:**
   - Prepare 500 µL of fresh 80% ethanol in nuclease-free water.
   - Spin down the sample and pellet the beads on a magnetic rack until the supernatant is clear.
   - Keeping the tube on the magnet, carefully remove the supernatant.
   - Wash the beads twice with 200 µL of 80% ethanol, ensuring the pellet remains undisturbed.
   - Remove any residual ethanol and allow the pellet to air-dry for ~30 seconds (avoiding over-drying to prevent cracking).
4. **Elution:**
   - Remove the tube from the magnetic rack and resuspend the pellet in 30 µL of nuclease-free water.
   - Incubate at room temperature for 2 minutes and return the tube to the magnetic rack.
   - Allow at least 1 minute for bead separation until the eluate is clear.
   - Carefully transfer 30 µL of eluate to a fresh 1.5 mL LoBind tube.

##### Adaptor Ligation and Purification

The entire purified DNA sample was used for the adaptor ligation step, following the recommended protocol (Table 2).

| Reagent | Volume (µL) |
|---|---|
| DNA sample from the previous step | 30 |
| Ligation Buffer (LNB) | 12.5 |
| NEB Next Quick T4 DNA Ligase | 5 |
| Ligation Adapter (LA) | 2.5 |

| | |
|---|---|
| Total | 50 |

***Table 3: Adaptor Ligation Reaction Setup***

1. **Adaptor Ligation Reaction:** The reaction was thoroughly mixed by gentle pipetting, briefly spun down, and incubated at room temperature for 10 minutes to facilitate ligation.
2. **AMPure XP Bead Purification:**
    - o The AMPure XP Beads were resuspended by vortexing.
    - o 20 µL of the resuspended AXP beads were added to the ligation reaction and mixed by flicking the tube.
    - o The mixture was incubated on a Hula mixer at room temperature for 5 minutes to allow DNA binding.
    - o The tube was spun down and placed on a magnetic rack until the beads formed a pellet. Once the supernatant was clear, it was carefully removed and discarded.

1. **Washing Steps:**
    - o 125 µL of Short Fragment Buffer (SFB) was added to the pellet, the tube was flicked to resuspend and then returned to the magnetic rack.
    - o Once the beads had pelleted, the supernatant was removed and discarded.
    - o The washing step was repeated to ensure complete removal of excess adaptors.
2. **Elution of DNA Library:**
    - o The tube was spun down and placed back on the magnet, with any residual supernatant removed.
    - o The pellet was allowed to air-dry for ~30 seconds, avoiding over-drying.
    - o The tube was removed from the magnetic rack, and the pellet was resuspended in 7 µL of Elution Buffer (EB).
    - o The mixture was incubated at room temperature for 10 minutes. For high molecular weight DNA, incubation at 37°C improved recovery.
    - o The tube was placed on the magnet until the eluate was clear and colorless (at least 1 minute).
    - o 7 µL of the purified DNA library was transferred to a clean 1.5 mL Eppendorf DNA LoBind tube, and the bead pellet was discarded.

3. **Final Quantification:**
    - The concentration of DNA sample was quantified before sequencing using 1ul of eluted DNA sample with Qubit fluorometer.

#### Final Preparation for Sequencing

Following purification, the sample was ready for sequencing. Before initiating the run, a flow cell quality check was performed to ensure optimal sequencing conditions. The sequencing reaction was set up as per ***Table 3***, and the Flow Cell ID was recorded. The MinION software was used to conduct a Flow Cell Check, where the number of active nanopores was assessed. The sample and pipettes were prepared for loading, ensuring proper handling to avoid contamination or air bubbles.

| Reagent | Volume (µL) |
|---|---|
| Sequencing Buffer (SB) | 15 |
| Library Beads (LIB) (mixed immediately before use) | 10 |
| DNA Library | 5 |

| | |
|---|---|
| Total | 30 |

***Table 4: Final Library Preparation Setup***

To flush the flow cell, 120 µL of flush buffer was loaded into the port. Before dispensing, the pipette plunger was unscrewed to remove any trapped air and expose a small drop of the sample. The pipette was held straight down into the exposed hole, ensuring a stable and secure fit before carefully dispensing the flush buffer by further unscrewing the plunger. Following the flush, 30 µL of the prepared DNA library was loaded using the same technique. Once the sample was successfully loaded, the sequencing run was initiated using MinION software, with the following settings applied like 20bp read length, POD5 detector and superaccurate basecalling. These settings ensured high-fidelity sequencing data, optimizing accuracy for downstream analysis.

#### Additional figures

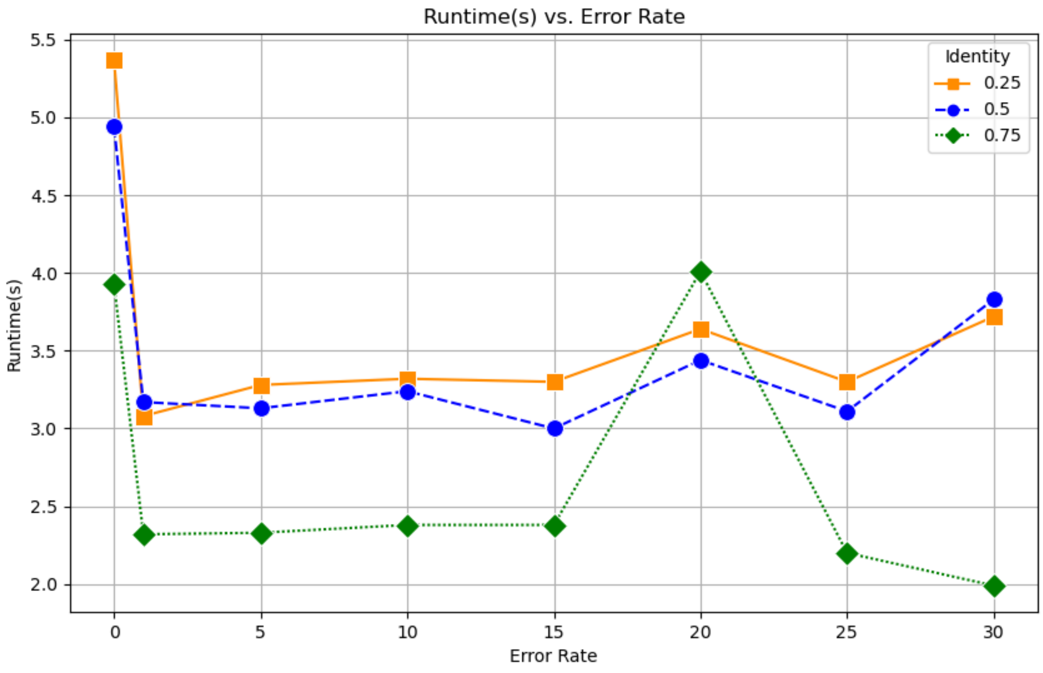


Supplementary Figure 1. Runtime in seconds of the cONcat algorithm for the SIM_0-SIM_30 datasets. Overall the runtime of cONcat is low due to the algorithm efficiently working on the sequencing data. A trend of runtime increase can be observed with lower identity settings with the runtime of SIM_20 for

0.75 exceeding the lower settings for the same error rate. This might be due to the computer having different workloads during the run of this dataset.

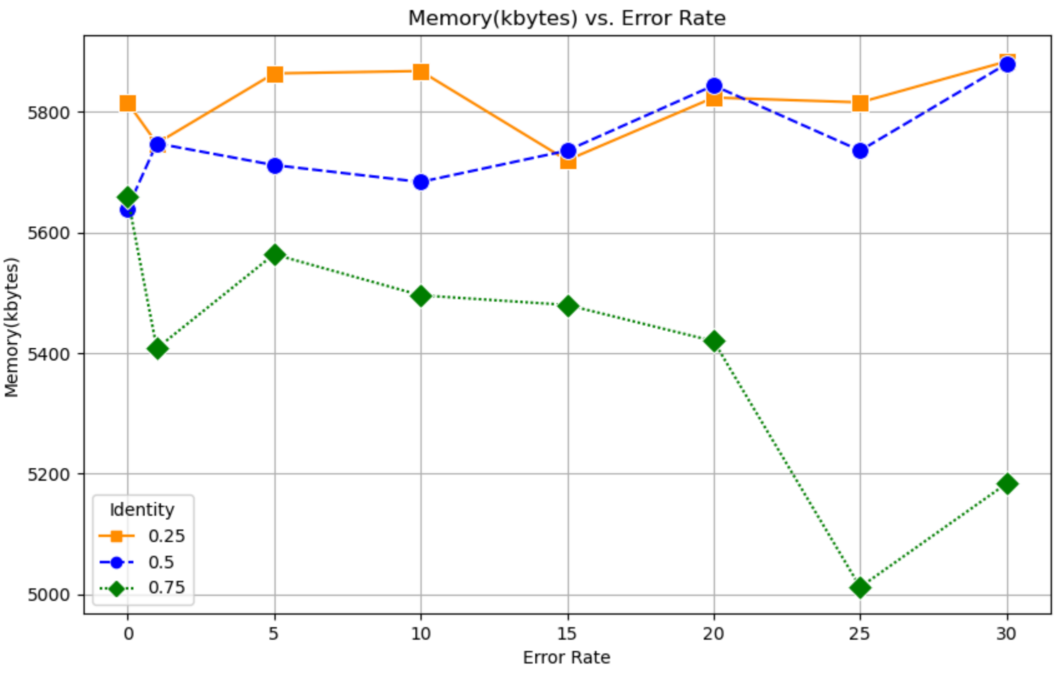


Supplementary Figure 2: Memory usage in kilobytes for the SIM_0-SIM_30 datasets of the cONcat algorithm. Overall the memory usage of cONcat is low due to the algorithm not having to store much data. The memory usage, however, shows a trend of slightly increasing with lower identity settings due to a higher amount of fragment hits being considered.
